# LAESI mass spectrometry imaging as a tool to differentiate the root metabolome of native and range-expanding plant species

**DOI:** 10.1101/322867

**Authors:** Purva Kulkarni, Rutger A. Wilschut, Koen J.F. Verhoeven, Wim H. van der Putten, Paolina Garbeva

## Abstract

Our understanding of chemical diversity in biological samples has greatly improved through recent advances in mass spectrometry (MS). MS-based-imaging (MSI) techniques have further enhanced this by providing spatial information on the distribution of metabolites and their relative abundance.

This study aims to employ laser-assisted electrospray ionization (LAESI) MSI as a tool to profile and compare the root metabolome of two pairs of native and range expanding plant species.

It has been proposed that successful range-expanding plant species, like introduced exotic invaders, have a novel, or a more diverse secondary chemistry. Although some tests have been made using aboveground plant materials, tests using root materials are rare. We tested the hypothesis that range-expanding plants possess more diverse root chemistries than native plant species.

To examine the root chemistry of the selected plant species, LAESI-MSI was performed in positive ion mode and data was acquired in a mass range of m/z 50-1200 with a spatial resolution of 100 µm. The acquired data was analyzed using in-house scripts, and differences in the spatial profiles were studied for discriminatory mass features.
The results revealed clear differences in the metabolite profiles amongst and within both pairs of congeneric plant species, in the form of distinct metabolic fingerprints. The use of ambient conditions and the fact that no sample preparation was required, established LAESI-MSI as an ideal technique for untargeted metabolomics and for direct correlation of the acquired data to the underlying metabolomic complexity present in intact plant samples.

## BACKGROUND

Detection of plant metabolites is extremely challenging, as there is no single-instrument platform available to effectively measure their overall coverage. During the last decade, mass spectrometry imaging (MSI) has emerged as a valuable tool, with numerous applications in the field of biological sciences. This analytical technique enables label-free, high-resolution spatial mapping of a large variety of biomolecules along with providing qualitative and quantitative chemical information, in a single experiment [1]. Identical to traditional mass spectrometry, during MSI it is important to ionize the sample to form ions suitable for mass analysis. Different ionization methods exist for MSI, however many of them require artificially altering the biochemical status of the system under study, for example by the use of a matrix, and are mainly operated under vacuum. Recently developed ambient ionization approaches such as laser ablation electrospray ionization (LAESI) allow direct analysis of biological samples in a matrix-free, native atmospheric condition with minimal to no sample preparation [2,3]. This opens up possibilities for in situ chemical analysis in biological systems.

LAESI-MSI is particularly tailored for biological samples that are rich in water content [4]. In this technique, the sample under investigation is mounted on a sample stage and is ablated using a focused mid-Infrared laser pulse, under atmospheric conditions. This ablation ejects a mixture of molecules, clusters, and particulate matter in microscopic volumes from the sample, in the form of a plume. The catapulted biomolecules then coalesce with charged droplets, produced by an electrospray to become ionized [5,6]. MSI using the LAESI ionization approach is realized by rastering the sample surface at pre-defined coordinates with a laser beam, where at each coordinate position the generated ions pass through the mass analyzer and a mass spectrum is recorded. LAESI-MSI has shown considerable success in revealing the lateral and cross-sectional distribution of primary and secondary metabolites for a range of plant related samples, along with providing chemical information from deeper parts of the tissue section [7]. LAESI equipped with a sharpened optical fiber tip has also been widely used to perform in situ metabolic profiling of single cells from plant and animal samples [8].

Here, we aim to demonstrate the potential of LAESI-MSI as an analytical technique for the direct metabolite profiling of plant samples. We applied LAESI-MSI in a comparative metabolic profiling study on two pairs of non-native, range-expanding plant species and congeneric native plant species. In response to recent climate warming, many plant species have expanded their range to higher latitudes and altitudes [9,10]. It is thought that plant secondary chemistry is an important factor determining the invasive success of exotic plant species. The novel chemistry of invasive exotic plant species may effectively control defenses against insect herbivores and other natural enemies [11]. Such ‘novel chemistry’ has been shown to potentially suppress native plant species directly through allelopathy [12] or indirectly through the suppression of the fungal mutualists of native plant species [13,14]. Moreover, due to this difference in chemistry, native generalist herbivores may perform less well on exotic plant species than on related native plant species [15], potentially leading to a reduced herbivore pressure on exotics compared to natives. The poor performance of generalist herbivores has also been linked to the high diversity of metabolites produced by exotic plant species compared to native plant species [16]. This suggests that chemically diverse plant species may be prone to become abundant when they are introduced in a new area where the local herbivores are poorly adapted to neutralize, or circumvent the novel defenses.

In this study, we use LAESI-MSI as a high-throughput tool for untargeted comparative metabolomics of intact plant roots of native and range expanding plant species. For this, we use two range-expanding plant species that are currently expanding in North-Western Europe, *Centaurea stoebe* L. and *Geranium pyrenaicum* Burm. f., and their respective congeneric native species *Centaurea jacea* L. and *Geranium molle* L. With this study, we demonstrate the suitability of LAESI-MSI for untargeted metabolomics profiling and we give insights in the potential chemical novelty of range-expanding plant species in comparison to congeneric related native plant species.

## DATA DESCRIPTION

In order to perform untargeted comparative root metabolomics using LAESI-MSI, two pairs of native plant species and their respective range-expanding species were selected. Intact root samples were collected from three biological replicates for the native species *Centaurea jacea* L. and *Geranium molle* L and their respective range expanding plant species *Centaurea stoebe* L. and *Geranium pyrenaicum* Burm. f. These twelve intact root samples were mounted on the sample stage one-by-one to perform LAESI-MSI in positive ion mode. The mass spectral data was acquired in a mass range of *m/z* 50-1200 from 105 pre-defined coordinate positions (spots) present on each sample replicate. The acquired data was first lock-mass corrected using an internal standard. Since all the 105 LAESI ablation spots for a single replicate were not present on the root sample, mass spectra arising from only 50 spots per replicate, that were visibly present on the root sample were selected manually. These extracted mass spectra were subjected to multiple data preprocessing steps. After performing peak detection on the preprocessed data, a mass feature matrix was generated. Multivariate data analysis was performed on the feature matrix to screen out significant differentially expressed metabolites amongst the samples.

## ANALYSES

### Untargeted metabolite profiling and multivariate analysis

Untargeted metabolomic studies are exploratory in nature and usually result in extremely large and multi-dimensional datasets. Analyses of such datasets using chemometric tools can hugely aid data interpretation.

The representative averaged preprocessed spectra for each replicate belonging to the different plant species show a clear distinction in mass spectra between the two plant genera (**Figure 1**). This distinction between *Centaurea* and *Geranium samples* was confirmed by unsupervised hierarchical clustering of the mass feature matrix (**Figure 2a**). Within the two genera, the different plant species were mostly clearly separated based on their chemical features, with the exception of one of the *C. stoebe* replicates (**Figure 2a**).

**Figure 1:**
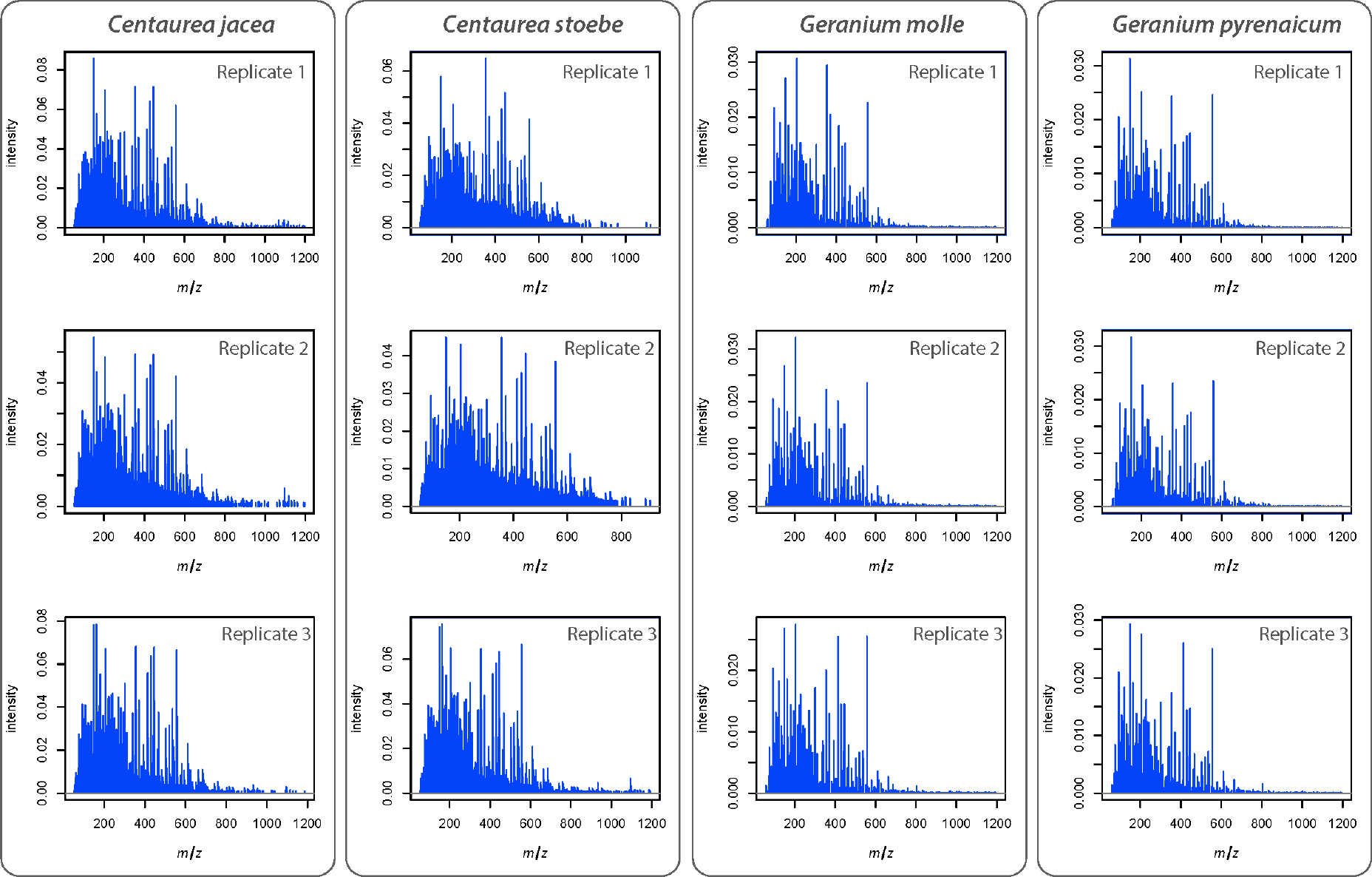
Metabolic profiling and comparison of LAESI-MS spectra from native and range expanding plant species. Each representative mass spectra is generated by averaging and preprocessing the signals acquired in positive ion mode, arising from the 50 ablation spots present on the imaged root sample for each replicate. The averaged preprocessed mass spectra are displayed for the three replicates of native species (*C. jacea* and *G. molle*) and the three replicates for range expanding plant species (*C. stoebe* L. and *G. pyrenaicum*).

Visual comparison of the representative mass spectrum for each sample group can be used to broadly study the differing metabolic profiles. To further examine these differences and similarities between the root metabolic profiles of the four plant species, we employed PCA. The first two selected principal component axes explain over 75% of cumulative variance amongst the samples (**Figure 2b**). Samples from different plant genera were strongly separated along the first PC-axis (~57%), whereas the separation along the second PC-axis (~18%) corresponded with within-genus variation (**Figure 2b**). Together with hierarchical clustering (**Figure 2a**), these results indicate a strong phylogenetic signal in root chemistry, as between-genus variation is considerably stronger than within-genus variation [17].

**Figure 2:**
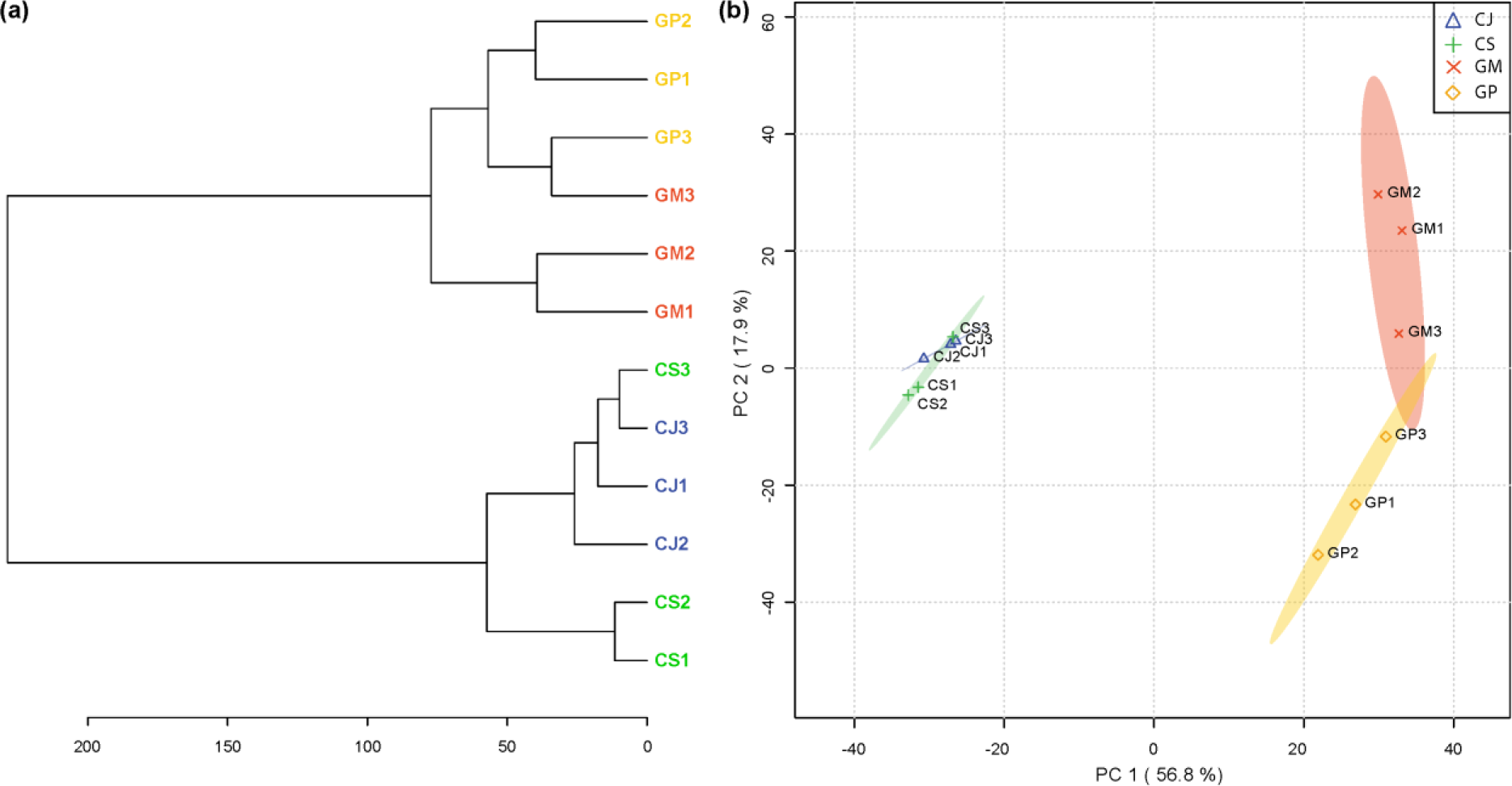
Heat map with dendrogram and Principal component analysis (PCA) score plot for the selected native and range expanding species. **(a)** Species clustering represented as a dendrogram (distance measure used is Euclidean and clustering algorithm is ward). Each node in the dendrogram corresponds to a single replicate belonging either to the range-expanding or to the congeneric native plant species. **(b)** The PCA score plot displays the total explained variance of >70 % for component 1 and component 2. Ovals represent 95% confidence intervals. Each oval represents a sample group and each point represents a single sample.

The number of mass features detected for each LAESI-MSI acquisition after performing data pre-processing and peak-detection clearly shows that there are more mass features detected for the replicates of *Centaurea* as compared to those of *Geranium* (**Table 1**). The two *Centaurea* species shared 314 metabolites, whereas 53 metabolites were unique to either one of the species (**Figure 3 a**). Interestingly, 49 of these metabolites were unique for range-expanding *C. stoebe*, whereas only 4 were unique for native *C. jacea*. In contrast, for native *G. molle* more unique metabolites were detected than in range-expanding *G. pyrenaicum* (**Figure 3b**). These results are in line with a previous study in which only root volatiles were examined [18] and indicate that range-expanding plants do not necessarily possess a more unique root chemistry than related natives.

**Table 1.** Overview of the number of metabolites detected in each sample replicate after preprocessing and peak detection of the acquired LAESI-MSI datasets.

| Replicate | *C. jacea* (CJ) | *C. stoebe* (CS) | *G. molle* (GM) | *G. pyrenaicum* (GP) |
| --- | --- | --- | --- | --- |
| 1 | 283 | 332 | 143 | 129 |
| 2 | 204 | 301 | 151 | 127 |
| 3 | 286 | 286 | 131 | 122 |

**Figure 3:**
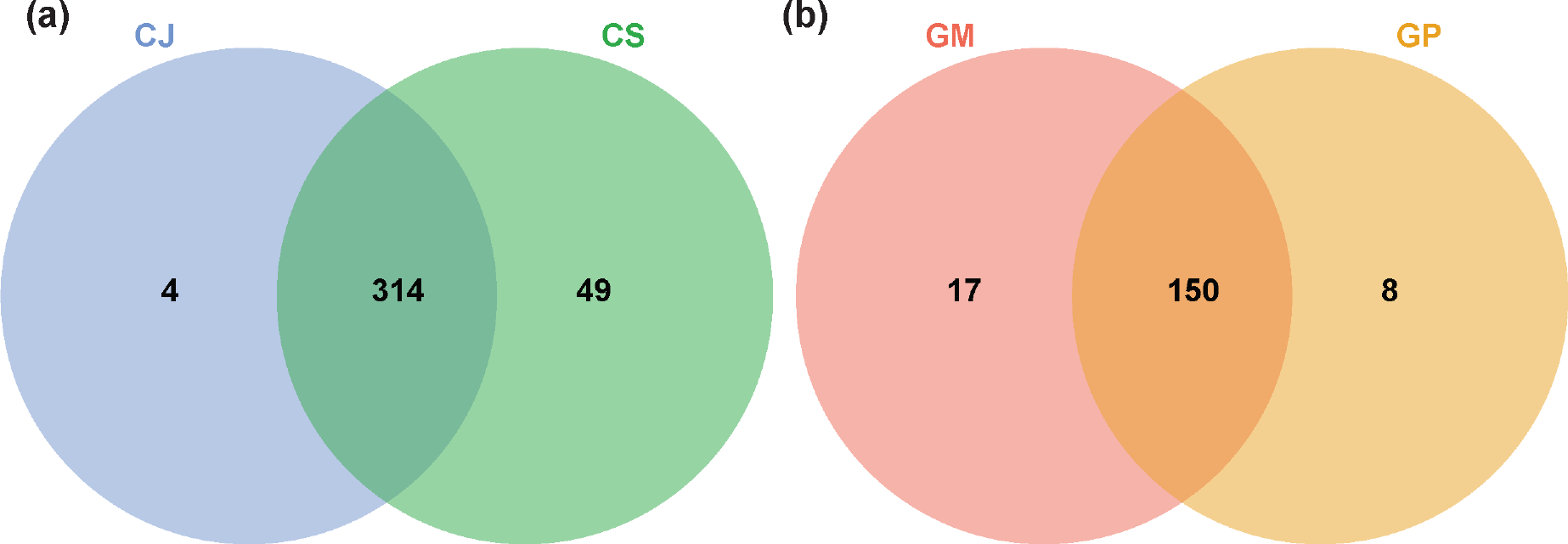
Venn diagram showing overlapping and unique metabolites associated with native and range expanding plant species. **(a)** Venn diagram for *C. jacea* (CJ) and *C. stoebe* (CS). **(b)** Venn diagram for *G. molle* (GM) and *G. pyrenaicum* (GP). To construct the Venn diagram, a single mass feature was considered even if it was present in only one replicate for a specific sample species.

In order to visualize the statistically significant metabolites for the two *Centaurea* species, a volcano plot was constructed (**Figure 4a**). As can be seen in **Figure 4a**, in total 367 metabolites were detected in genus *Centaurea*. Within this, 10 mass features (shown in green) that are located in upper right quadrant of the plot, indicate that their concentration is significantly higher in native species *C. jacea* than in range expanding species *C. stoebe*. The 5 mass features (shown in red) that are observed in the upper left quadrant indicate that their concentration is significantly lower in native species *C. jacea* than in range expanding species *C. stoebe*. To examine the differences in metabolite concentrations for the *C. jacea* and *C. stoebe* pair, box-and-whisker plots were realized for four statistically significant metabolites chosen based on the volcano plot (**Figure 4b**). As can be seen in box-and-whisker plots and the ion intensity maps, *m/z* 84.9607, *m/z* 159.0520 and *m/z* 557.290 are highly abundant in native species *C. jacea*, whereas *m/z* 272.9550 are highly abundant in range-expanding species *C. stoebe*. Additionally, the corresponding ion intensity maps for these metabolites were also generated to visualize the changes on the spatial level in the imaged roots. The ion intensity maps can be seen alongside the box-and-whisker plots in **Figure 4b**. Each ion map is plotted on the same color scale (depicted below the ion maps) ranging from 0 (blue meaning least intense) to 1 (red meaning most intense), to allow comparison of relative ion intensity between images.

**Figure 4:**
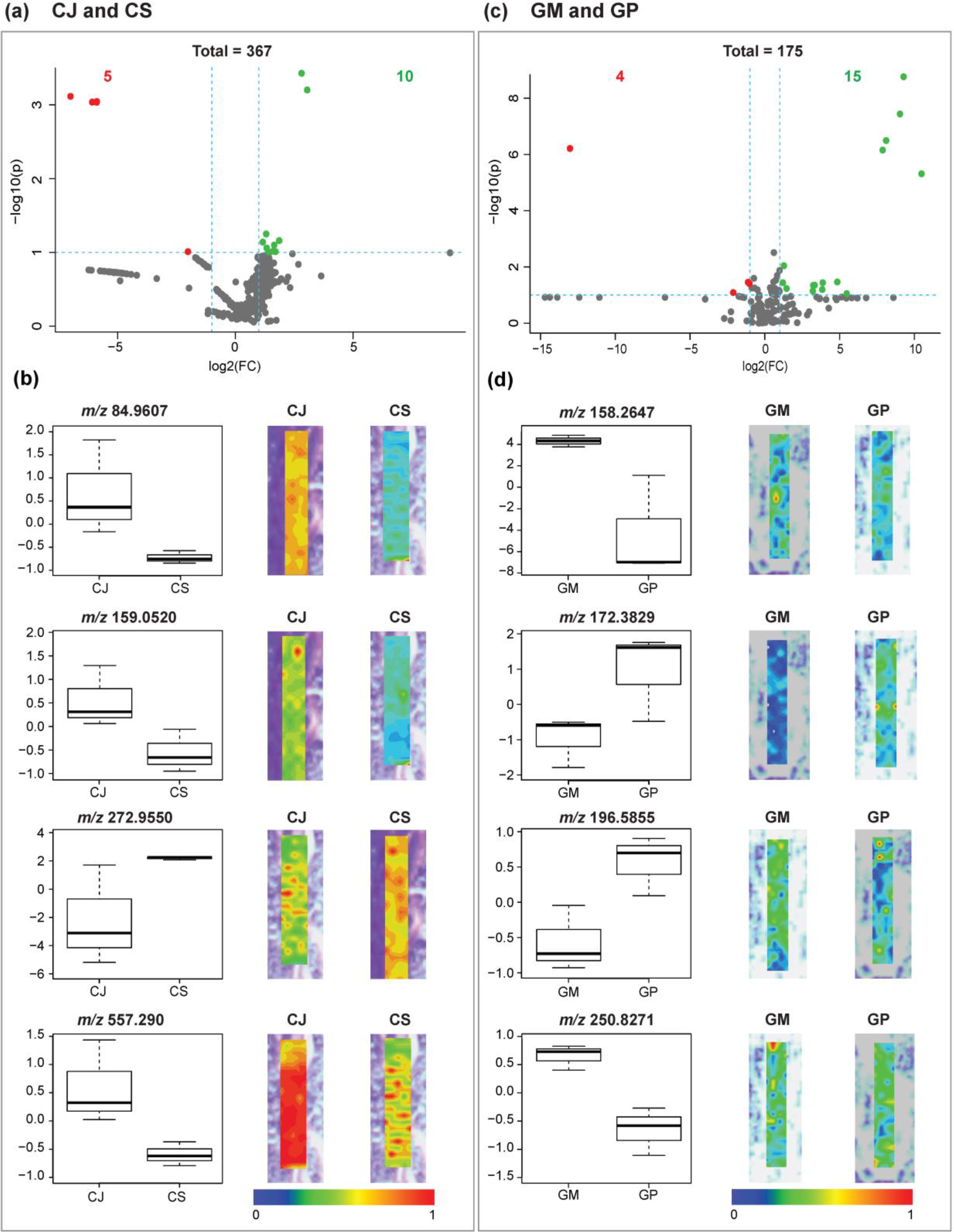
Volcano plots and box plots to demonstrate metabolite concentration differences observed in native and range expanding plant species. **(a)** Volcano plot for *C. jacea* (CJ) vs. *C. stoebe* (CS). **(b)** Volcano plot for *G. molle* (GM) vs. *G. pyrenaicum* (GP). Each point in the volcano plot represents one metabolite. Significant metabolites were calculated with a fold change (FC) threshold of 2 on the x-axis and a t-tests threshold of 0.1 on the y-axis. The red and the green dots indicate statistically significant metabolites, and the gray dots below the FC threshold line represent statistically non-significant metabolites. The vertical FC threshold lines indicates an increase or decrease in concentration of metabolites. Negative log2 (FC) values indicated in red represent lower concentrations in native than in range expanding species; positive values indicated in green represent higher concentrations of metabolites in native than in range expanding species. The box plots for the detected metabolites and their corresponding ion intensity maps below each volcano plot display the localization of the selected metabolites that are significantly different in the respective native and range expanding species. The signal intensity in the ion intensity maps are represented in rainbow color scale, in a mass window of ±1 mDa.

Similar analysis was performed for the two *Geranium* species (**Figure 4c**). For this pair, in total 175 metabolites were detected. Within these, 15 mass features (shown in green) that are located in the upper right quadrant of the plot, which indicates that their concentrations are significantly higher in native species *G. molle* than in range expanding species *G. pyrenaicum*. The 4 mass features (shown in red) that are observed in the upper left quadrant indicates that their concentration is significantly lower in native species *G. molle* than in range expanding species *G. pyrenaicum*. The box-and-whisker plots for the four statistically significant metabolites selected from the volcano plot for the pair *G. molle* and *G. pyrenaicum* are shown in **Figure 4d**. The ion intensity maps for these statistically significant metabolites are shown alongside box-and-whisker plots. As can be seen, *m/z* 158.2647 and *m/z* 250.8271 show high abundance in native species *G. molle*, whereas *m/z* 172.3829 and *m/z* 196.5855 display high abundance in range expanding species *G. pyrenaicum*. All significant metabolites detected for *Centaurea* and *Geranium samples* are listed in **Supplementary Table 1**.

## DISCUSSION

In this study, we demonstrated the utility of the ambient ionization ability of LAESI coupled with MSI to explore the chemical differences in the root metabolome between two pairs of native and range expanding plant species. This high-throughput technology provided an *in situ* analysis method capable of revealing differentially produced metabolites linked to each group. We detected clear differences in root chemical profiles within both pairs of range-expanding plant species and congeneric natives using untargeted LAESI-MSI approach. Interestingly, the range-expanding plant species *Centaurea stoebe* showed a strongly unique root chemistry, which also may have enabled this species to become invasive in its introduced range in North America [15,19].

Furthermore we demonstrated that LAESI-MSI can help to spatially elucidate the metabolite composition of the intact roots with minimal to no sample preparation. Our demonstration did not involve an exhaustive region-specific spatial analysis of the roots, but rather a ‘proof of concept’ by lateral profiling of the root samples. This allowed us to establish that LAESI-MSI of whole-root sections could reveal information on location-specific metabolite distribution without the need for any sample preparation. These results can help to reveal the role of single metabolites based on their location within the roots.

Overall, our results illustrate the feasibility of LAESI-MSI as a high-throughput technique for the detection and localization of metabolites from intact plant samples and gaining spatial information without the need for extensive sample preparation. The potential applications of this work could lead to rapid phenotyping of plant tissues as well as comparative untargeted metabolomics of different plant parts, a topic of considerable recent interest for plant research.

## METHODS

### Plant species and root collection

The seeds used for all four plant species originated from natural populations in natural areas in The Netherlands, where the range expanders are immigrating. Seeds of *G. molle* and *C. stoebe* were collected directly from the field. For *C. jacea*, seeds were collected from plants growing in an experimental garden, whereas the mother plants were germinated from field-collected seeds. Seed production company Cruydt-hoeck (Groningen, The Netherlands), that grows plants originating from field-collected seeds, delivered the seeds for *G. pyrenaicum*. For all plant species, the seeds were surface-sterilized by washing for 3 min in a 10% bleach solution, followed by rinsing with demineralized water, after which they were germinated on glass beads. After 20 days, the seedlings were collected for LAESI analysis.

### LAESI mass spectrometry imaging

The LAESI-MSI of intact roots collected from the seedlings was carried out on a Protea Biosciences DP-1000 LAESI system (Protea Bioscience Inc., Morgantown) coupled to a Waters model Synapt G2S (Waters Corporation) mass spectrometer. The LAESI system was equipped with a 2940-nm mid-infrared laser yielding a spot size of 200 µm. The laser was set to fire 10 times per x-y location (spot) at a frequency of 10 Hz and 100% output energy. The system was set to shoot at 105 locations per plant root (grid of 21 × 5 positions). A syringe pump was delivering the solvent mixture of methanol/water/formic-acid (50:50:0.1% v/v) at 2 µL/min to a PicoTip (5cm × 100 µm diameter) stainless steel nanospray emitter operating in positive ion mode at 3800 V. The LAESI was operated using LAESI Desktop Software V2.0.1.3 (Protea Biosciences Inc.). The Time of Flight (TOF) mass analyzer of the Synapt G2S was operated in V-reflectron mode at a mass resolution of 18.000 to 20.000. The source temperature was 150 ±C, and the sampling cone voltage was 30 V. The data was acquired in a mass range of *m/z* 50 to 1200. The acquired MS data was lock mass corrected post data acquisition using leucine encephalin (C_28_H_37_N_5_O_7_, *m/z* = 556.2771), which was added in the spray as an internal standard.

### Data processing, peak-detection and chemometrics

All the acquired Waters ․raw data files were first pre-processed to remove noise and to make the data comparable. Since the root samples used in this study were tiny, many LAESI ablation spots constituted the background on which the root samples were placed. In order to avoid including the mass spectra purely consisting of spectral signals from the background, 50 ablation spots per sample replicate, present on the root section were selected manually. The selected ablation spots for every sample replicate are displayed in **Supplementary Figure 1**. The mass spectra arising from the spots colored in green are included in the study whereas those in red have been excluded.

The spectra from all the 50 selected spots for each replicate were averaged. Processing of these mass spectra involved multiple steps. An overview of the data processing steps applied is provided in **Figure 5**. First, square root transformation was applied to overcome the dependency of variance on the mean. Then, baseline correction was performed to enhance the contrast of peaks to the baseline. For better comparison of intensity values and to remove small batch effects, Total-Ion-Current (TIC)-based normalization was applied. This was followed by spectral alignment and peak detection to extract a list of significant mass features for each sample replicate. In the end, a mass feature matrix was generated with sample replicates in columns and mass features in rows. This feature matrix was used to perform chemometric analysis. The preprocessing and peak-detection steps were applied using R scripts developed in-house and the functions available within the MALDIquant R package [20].

**Figure 5:**
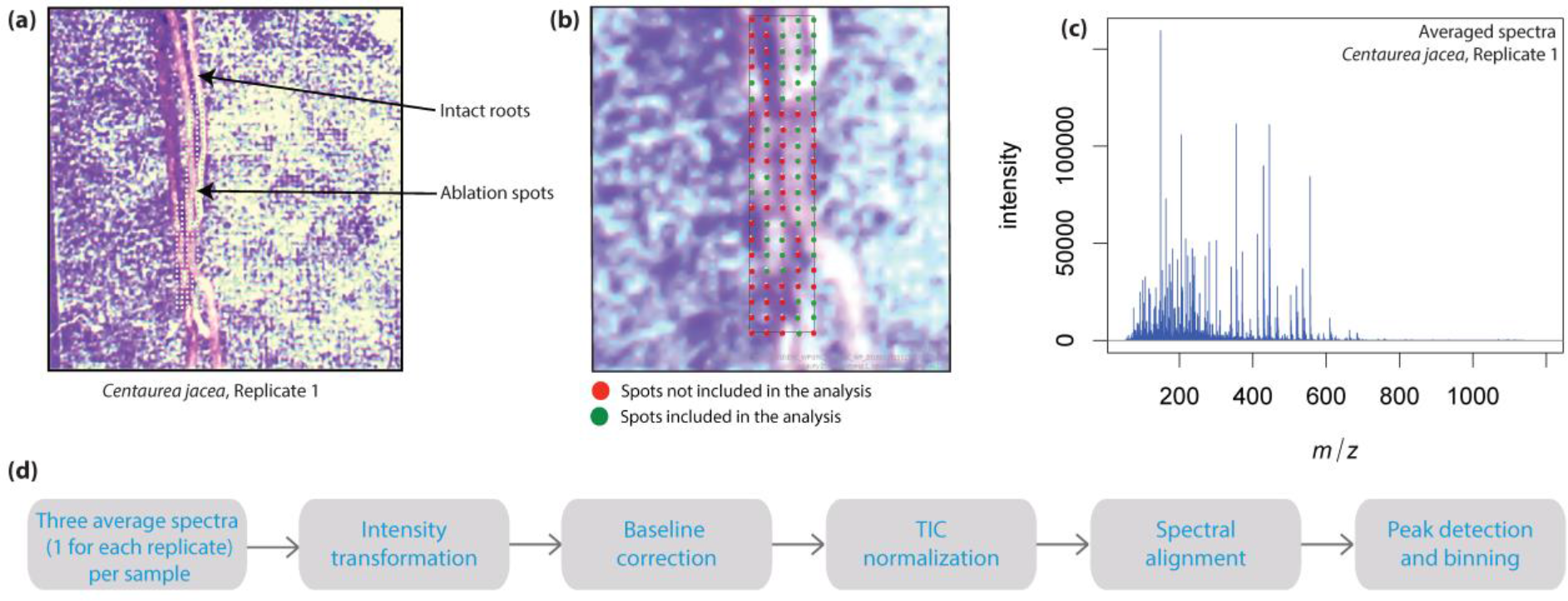
Data preparation and processing steps applied post acquisition. **(a)** Optical image for the intact roots of a single replicate of *C. jacea* with labeled ablation spots. **(b)** Ablation spots present on the root selected (in green) for further analysis. **(c)** Averaged spectra acquired from all the 50 selected spots per replicate. **(d)** Data pre-processing and peak-detection steps applied to all spectra for a sample.

To perform multivariate analysis, the feature matrix was imported into Metaboanalyst 3.0 [21]. Principal component analysis (PCA), was initially applied to visualize the intrinsic spectral differences in the non-native, range-expanding plant species and congeneric native plant species. In order to get an overview of the differences amongst the samples, a dendrogram showing clustering of the sample replicates was generated, using the Euclidean distance measure and the Ward’s clustering algorithm.

To visualize the number of differential metabolites in in non-native, range-expanding plant species and congeneric native plant species, a pairwise comparative analysis was performed. To graphically illustrate these differences volcano plots were generated. Metabolites with a fold change (FC) threshold of 2 on the x-axis and a t-tests threshold (p-value) of 0.1 on the y-axis were considered significant. Box plots for selected significant metabolites were created to display changes in the concentration of native and range-expanding species. Corresponding accurate ion intensity maps (±1 ppm) displaying spatial distribution for these selected mass features were created using the ProteaPlot software V2.0.1.3 (Protea Biosciences Inc., Morgantown, WV). Venn diagrams were drawn using the jvenn tool [22] to plot the number of shared and unique metabolites for each pair of samples.

## AVAILABILITY OF SOURCE CODE AND REQUIREMENTS

Project name: LAESI-MSI-Root-Metabolomics

Project home page: https://github.com/purvakulkarni7/LAESI-MSI-Root-Metabolomics

Operating system(s): platform independent

Programming language: R

Other requirements: R (≥ 3.2.0), MALDIquant package, MALDIquantForeign package

License: GNU General Public License version 2.0 (GPLv2).

Any restrictions to use by non-academics: none

## ABBREVIATIONS

MS: mass spectrometry; MSI: mass spectrometry imaging; LAESI: laser-assisted electrospray ionization; CJ: *Centaurea jacea* L.; CS: *Centaurea stoebe* L.; GM: *Geranium molle* L.; GP: *Geranium pyrenaicum* Burm.; *m/z*: mass by charge; GC: gas chromatography; TOF: Time-of-flight; PCA: principal component analysis; PC: principal component; FC: fold change.

## ACKNOWLEDGEMENTS

We thank Frank Claassen from the department of Agrotechnology and Food Sciences at the Wageningen University for assistance with LAESI-MSI measurement and Julio Pereira da Silva for help with the experimental preparation. Purva Kulkarni is supported by the strategic project fund from the Netherlands Institute of Ecology (NIOO-KNAW). Rutger A. Wilschut and Wim H. van der Putten are supported by the ERC advanced grant ERC-Adv 26055290. This is publication XXX of the Netherlands Institute of Ecology (NIOO-KNAW).

## AUTHOR CONTRIBUTIONS

P.G., K.J.F.V. and R.A.W. devised the project. P.G. and R.A.W. oversaw the sample collection and the data acquisition. P.K. planned and performed the bioinformatics analysis, interpretation of results and prepared the figures. P.K., R.A.W. and P.G. wrote the manuscript. W.H.v.d.P., P.G., K.J.F.V. and R.A.W. provided their comment and contributed to substantial revision of the manuscript.

---

**Supplementary Figure 1:**
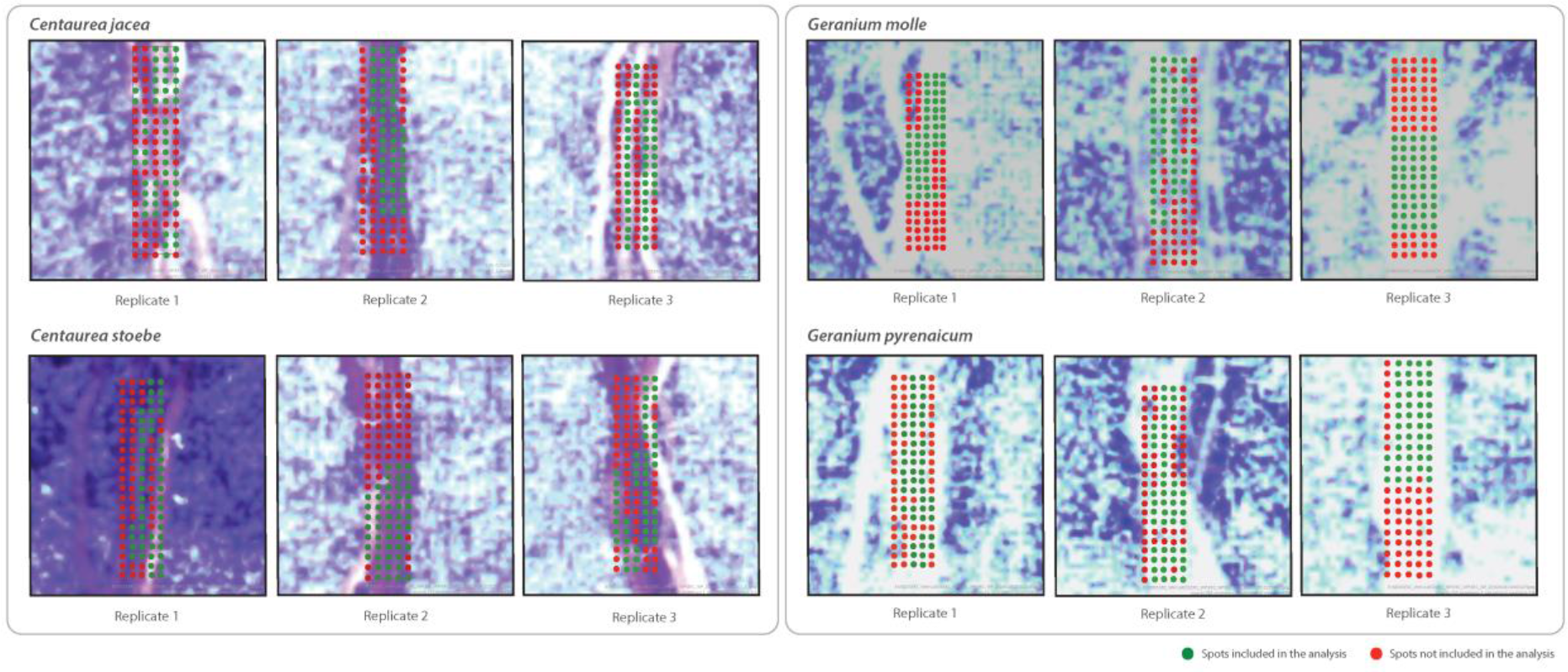
Ablation spots present on the imaged root samples selected for further analysis. A set of 50 ablation spots for each replicate of the native and range expanding species was selected. The spots selected for further analysis are shown in green. These are present on the root sample that has been imaged. The spots that are not selected for further analysis are displayed in red. These may or may not arise from the imaged root samples.

**Supplementary Table 1:** Significant metabolites and their respective fold change (log_2_(FC)) and *p* values (−log_10_(p)) for native and range expanding pairs *C. jacea* (CJ) vs. *C. stoebe* (CS) and *G. molle* (GM) vs. *G. pyrenaicum* (GP).

| CJ vs. CS |  |  | GM vs. GP |  |  |
| --- | --- | --- | --- | --- | --- |
| *m/z* | $\log_2(\text{FC})$ | $-\log_{10}(p)$ | *m/z* | $\log_2(\text{FC})$ | $-\log_{10}(p)$ |
| 892.2366 | 2.8154 | 3.4282 | 887.111 | 9.2804 | 8.7636 |
| 837.1989 | 3.0537 | 3.2022 | 885.1207 | 9.039 | 7.44 |
| 245.094 | -7.0019 | 3.1155 | 980.8683 | 8.1151 | 6.4941 |
| 270.964 | -5.8824 | 3.0457 | 486.52534 | -13.033 | 6.2126 |
| 213.12 | -6.0788 | 3.037 | 1096.874 | 7.8812 | 6.1562 |
| 352.952 | -5.8897 | 3.0347 | 492.5648 | 10.482 | 5.3132 |
| 557.2904 | 1.3107 | 1.2492 | 250.8271 | 1.2766 | 2.0428 |
| 136.0762 | 1.8652 | 1.1594 | 158.2647 | 4.8462 | 1.4655 |
| 159.0517 | 1.1667 | 1.1396 | 196.58554 | -1.1204 | 1.455 |
| 84.96074 | 1.65 | 1.0988 | 252.3956 | 1.2045 | 1.4386 |
| 536.1757 | 1.3311 | 1.0584 | 1142.875 | 3.8741 | 1.4379 |
| 99.00528 | 1.6044 | 1.0152 | 64.50864 | -1.0269 | 1.4038 |
| 87.0236 | 1.7072 | 1.0096 | 922.8092 | 3.3317 | 1.3522 |
| 272.955 | -2.013 | 1.0087 | 1024.339 | 3.2395 | 1.3445 |
| 59.02047 | 1.4073 | 1.0044 | 637.3655 | 1.4521 | 1.2323 |
|  |  |  | 187.6822 | 3.8414 | 1.1946 |
|  |  |  | 92.45683 | 3.2114 | 1.1408 |
|  |  |  | 172.38286 | -2.1105 | 1.0878 |
|  |  |  | 859.1226 | 5.4774 | 1.0524 |

